# Reproducible integration of multiple sequencing datasets to form high-confidence SNP, indel, and reference calls for five human genome reference materials

**DOI:** 10.1101/281006

**Authors:** Justin M. Zook, Jennifer McDaniel, Hemang Parikh, Haynes Heaton, Sean A. Irvine, Len Trigg, Rebecca Truty, Cory Y. McLean, Francisco M. De La Vega, Chunlin Xiao, Stephen Sherry, Marc Salit

## Abstract

Benchmark small variant calls from the Genome in a Bottle Consortium (GIAB) for the CEPH/HapMap genome NA12878 (HG001) have been used extensively for developing, optimizing, and demonstrating performance of sequencing and bioinformatics methods. Here, we develop a reproducible, cloud-based pipeline to integrate multiple sequencing datasets and form benchmark calls, enabling application to arbitrary human genomes. We use these reproducible methods to form high-confidence calls with respect to GRCh37 and GRCh38 for HG001 and 4 additional broadly-consented genomes from the Personal Genome Project that are available as NIST Reference Materials. These new genomes’ broad, open consent with few restrictions on availability of samples and data is enabling a uniquely diverse array of applications. Our new methods produce 17% more high-confidence SNPs, 176% more indels, and 12% larger regions than our previously published calls. To demonstrate that these calls can be used for accurate benchmarking, we compare other high-quality callsets to ours (e.g., Illumina Platinum Genomes), and we demonstrate that the majority of discordant calls are errors in the other callsets, We also highlight challenges in interpreting performance metrics when benchmarking against imperfect high-confidence calls. We show that benchmarking tools from the Global Alliance for Genomics and Health can be used with our calls to stratify performance metrics by variant type and genome context and elucidate strengths and weaknesses of a method.

## Introduction

As genome sequencing is increasingly used in clinical applications, high-quality variant and reference genotype calls for a small number of genomes consented for open dissemination are essential as part of benchmarking variant call accuracy. The Genome in a Bottle Consortium (GIAB) was formed as an open science project for authoritative characterization of benchmark genomes by integrating multiple technologies and bioinformatics methods. We previously described methods to form high-confidence SNP, indel, and reference genotype calls for the pilot GIAB genome, NIST Reference Material (RM) 8398 (which is NIST sample HG001, from the same cell lines as the Coriell DNA NA12878).^1^ These benchmark calls have been used as part of optimization and analytical validation of clinical sequencing,^2,3^ comparisons of bioinformatics tools,^4^ and optimization, development, and demonstration of new technologies.^5^

In this work, we build on our previous integration methods to enable the development of highly accurate, reproducible benchmark genotype calls from any genome with multiple datasets from different sequencing methods (Figure 1). We first develop a new version of more comprehensive and accurate integrated SNP, small indel, and homozygous reference calls for HG001. Phased pedigree-based callsets (from the Illumina Platinum Genomes Project^6^ and by Cleary et al.^7^) provide an orthogonal confirmation of variants by phasing variants in HG001, her 11 offspring and their father, and then verifying that the variants and haplotypes are inherited as expected from Mendelian segregation. We compare these pedigree-based callsets to ours following best practices established by the Global Alliance for Genomics and Health Benchmarking Team,^8^ and we manually curate a subset of differences between the callsets to understand the sources of disagreement.

**Figure 1:**
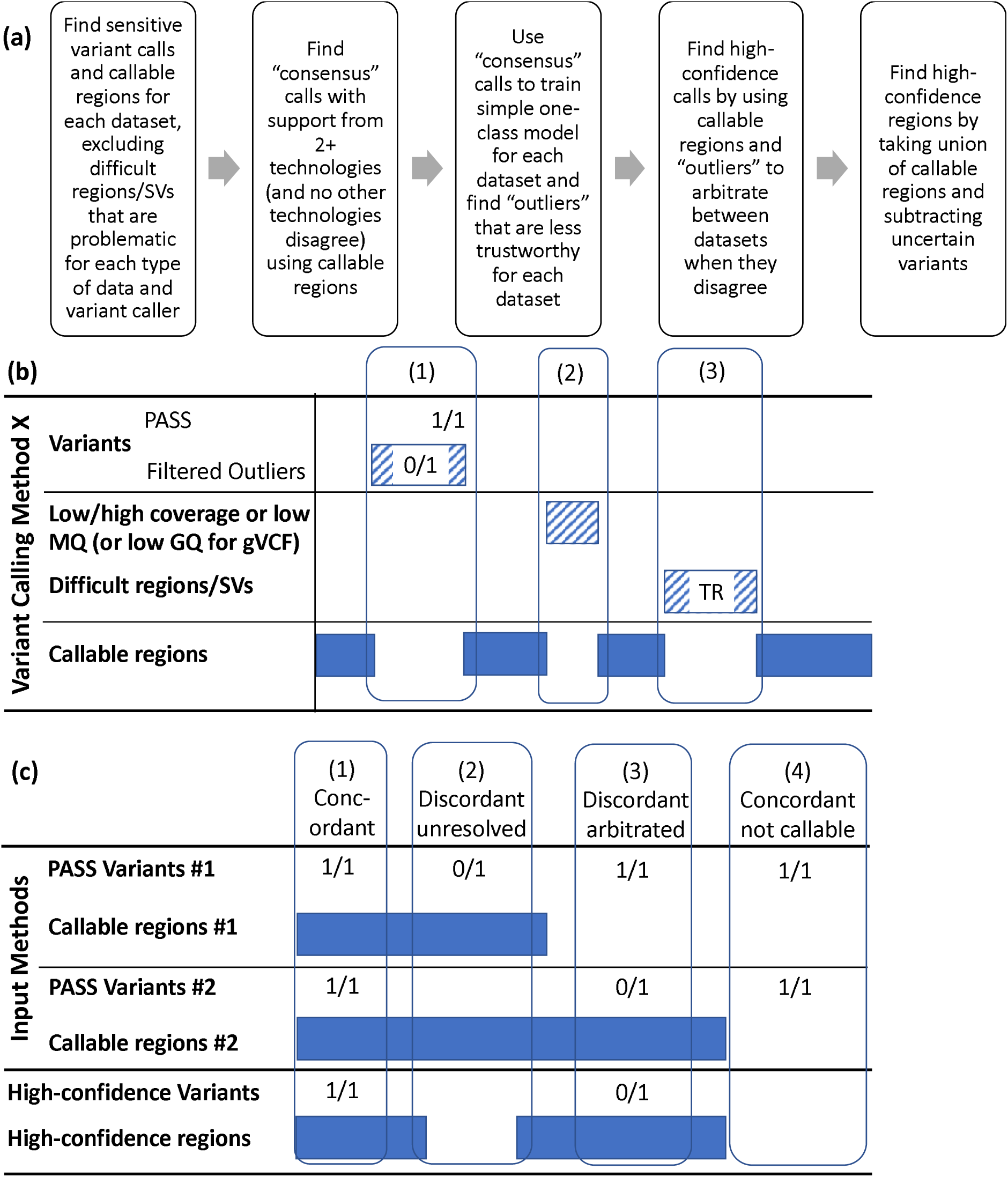
Overview of integration process and examples of the arbitration process used to integrate multiple technologies and callsets. The arbitration process has two cycles. The first cycle ignores “Filtered Outliers”. Calls that are supported by at least 2 technologies in the first cycle are used to train a model that identifies variants from each callset with any annotation value that is an “outlier” compared to these 2-technology calls. In the second cycle, the outlier variants and surrounding 50bp are excluded from the callable regions for that callset. (b) For each variant calling method, we delineate callable regions by subtracting (1) regions around filtered/outlier variants, (2) regions with low coverage or mapping quality (MQ), and (3) “difficult regions” prone to systematic miscalling or missing variants for the particular method. For callsets in gVCF format, we exclude homozygous reference regions and variants with genotype quality (GQ) < 60. Difficult regions include different categories of tandem repeats (TR) and segmental duplications. (c) Four arbitration examples with two arbitrary input methods. (1) Both methods have the same genotype and variant and it is in their callable regions, so the variant and region are high-confidence. (2) Method 1 calls a heterozygous variant and Method 2 implies homozygous reference, and it is in both methods’ callable regions, so the discordant variant is not high-confidence and 50bp on each side is excluded from the high-confidence regions. (3) The methods have discordant genotypes, but the site is only inside Method 2’s callable regions, so the heterozygous genotype from Method 2 is trusted and is included in the high-confidence regions. (4) The two methods’ calls are identical, but they are outside both methods’ callable regions, so the site is excluded from high-confidence variants and regions.

We apply our methods to four broadly consented GIAB genomes from the Personal Genome Project (PGP),^9^ an Ashkenazim Jewish (AJ) mother-father-son trio whose DNA is in NIST RMs 8391 and 8392 and the son of a Chinese trio whose DNA is NIST RM 8393. We use extensive data for these genomes that GIAB has generated using a variety of technologies.^10^ We also use these new methods to form similar high-confidence calls with respect to GRCh38. In the future, these methods can be used to characterize the parents of the Chinese trio, as well as other genomes characterized by multiple technologies.

Our extensively characterized genomes from PGP have an open, broad consent and broadly available data and cell lines, as well as ability to re-contact for additional types of samples.^9^ They are an enduring resource uniquely suitable for diverse research and commercial applications. A variety of products using these genomes is already available, including induced pluripotent stem cells (iPSCs), cell line mixtures, cell line DNA with synthetic DNA spike-ins with mutations of clinical interest, formalin-fixed paraffin-embedded (FFPE) cells, and circulating tumor DNA mimics.

Our work on broadly consented and available samples complements recent work to characterize samples with more restricted availability. These more restricted samples have been characterized for variants and regions not characterized with high-confidence by GIAB in this manuscript (e.g., Platinum Genomes pedigree analysis,^6^ HuRef analysis using Sanger sequencing,^11^ integration of multiple SV calling methods on HS1011,^12^ and synthetic diploid using long-read sequencing of hyditaform moles that are not broadly available^13^).

Below, we describe our methods used to integrate multiple short and linked read data types to form high-confidence SNV, small indel, and homozygous reference genotypes for these genomes. We systematically evaluate the resulting genotypes to demonstrate that they can be used to identify false positive and false negative calls with confidence.

## Results

### Design of high-confidence calls

Our goal is to design *reproducible, robust, and flexible* methods to produce high-confidence *variant* and *genotype* calls (including homozygous reference regions) that, when any sequencing method is compared to our high-confidence calls requiring *stringent matching of alleles and genotypes*, the majority of *discordant calls in the high-confidence regions* (i.e., FPs and FNs) *should be* attributable to *errors in the sequencing method*. We develop a modular, cloud-based data integration methods to make a reproducible, robust, and flexible pipeline, enabling diverse data types to be integrated for each genome. We produce high-confidence variant calls and regions by integrating methods and technologies with different strengths and limitations, and using evidence of potential bias to arbitrate when methods have differing results (Fig. 1). Finally, we evaluate the utility of the high-confidence variants and regions by comparing high-quality callsets to the high-confidence calls and manually curating discordant calls to ensure most are errors in the other high-quality callsets.

### New high-confidence calls are more comprehensive and accurate

Table 1 shows the evolution of the GIAB/NIST high-confidence calls since our previous publication. The fraction of non-N bases in the GRCh37 reference covered has increased from 77.4% to 90.8%, and the numbers of high-confidence SNPs and indels have increased by 17% and 176%, respectively. The fraction of GRCh37 RefSeq coding regions covered has increased from 73.9% to 89.9%. The gains in high-confidence regions and SNPs are a result of a less conservative SV bed file, as well as a new approach that excludes different types of difficult regions for different input callsets depending on read length, error profiles, and analysis methods. The larger gains in indels result from newer input callsets that have more accurate and sensitive indel calls, as well as newer integration methods that take into account that some input callsets are not sensitive to larger indels. The slight decrease in high-confidence calls from v3.3.1 to v3.3.2 resulted from fixing a problem in how v3.3.1 was calculating callable regions from the GATK gvcf. The smaller number of high-confidence bases and variants in GRCh38 appears to be largely due to a problem in the self-chain segmental duplication bed file, which causes some large regions to be erroneously excluded from the high-confidence regions. We do not expect this to affect recall and precision but will be fixed in future versions.

**Table 1:** Summary of statistics of GIAB high-confidence (highconf) calls and regions from HG001 from v2.18 to v3.3.2 and their comparison to Illumina Platinum Genomes 2016-v1.0 calls (PG). Note that PG bed files were contracted by 50bp to minimize partial complex variant calls in the PG calls.

| Integration Version | v 2.18 | v 2.19 | v. 3.2 | v. 3.2.2 | v 3.3 | v 3.3.1 | v 3.3.2 | v 3.3.1 | v 3.3.2 |
| --- | --- | --- | --- | --- | --- | --- | --- | --- | --- |
| Reference | GRCh37 | GRCh37 | GRCh37 | GRCh37 | GRCh37 | GRCh37 | GRCh37 | GRCh38 | GRCh38 |
| Integration Date | Sep 2014 | Apr 2015 | May 2016 | June 2016 | Aug 2016 | Oct 2016 | Nov 2016 | Oct 2016 | Nov 2016 |
| Number of bases in highconf regions (chromosomes 1-22 + X) | 2.20 Gb | 2.22 Gb | 2.54 Gb | 2.53 Gb | 2.57 Gb | 2.58 Gb | 2.58 Gb | 2.45 Gb | 2.44 Gb |
| Fraction of non-N bases covered in chromosomes 1-22 + X) | 77.4% | 78.1% | 89.6% | 89.2% | 90.5% | 91.1% | 90.8% | 84.2% | 83.8% |
| Fraction of RefSeq coding sequence covered | 73.9% | 74.0% | 87.8% | 87.9% | 89.9% | 90.0% | 89.9% | 83.4% | 83.3% |
| Total number of calls in highconf regions | 2915731 | 3153247 | 3433656 | 3512990 | 3566076 | 3746191 | 3691156 | 3617168 | 3542487 |
| SNPs | 2741359 | 2787291 | 3084406 | 3154259 | 3191811 | 3221456 | 3209315 | 3058368 | 3042789 |
| Insertions | 86204 | 172671 | 171866 | 176511 | 171715 | 243856 | 225097 | 269331 | 241176 |
| Deletions | 87161 | 189932 | 169389 | 173976 | 189807 | 266386 | 245552 | 275041 | 247178 |
| Block Substitutions | 1005 | 2532 | 7476 | 7716 | 10364 | 13332 | 11192 | 13976 | 11344 |
| Transition/Transversion ratio for SNVs | 2.12 | 2.12 | 2.14 | 2.14 | 2.11 | 2.10 | 2.10 | 2.10 | 2.11 |
| Fraction phased globally or locally | 0.0% | 0.3% | 3.9% | 3.9% | 8.8% | 99.0% | 99.6% | 98.5% | 99.5% |
| **High-confidence Comparisons to Platinum Genomes (PG) calls** |  |  |  |  |  |  |  |  |  |
| Number of highconf calls concordant with PG in both PG and GIAB beds | 2825803 | 3030703 | 3312580 | 3391783 | 3441361 | 3550914 | 3529641 | 3459674 | 3431752 |
| Number of PG-only calls in both beds | 194 | 404 | 81 | 52 | 60 | 67 | 61 | 202 | 180 |
| Number of GIAB-only calls in both beds | 49 | 87 | 56 | 57 | 40 | 50 | 47 | 105 | 94 |
| Number of PG-only calls: Total | 1223697 | 1018795 | 274671 | 138894 | 550982 | 445563 | 469202 | 659870 | 690887 |
| After excluding PG bed | 605142 | 493694 | 118810 | 22224 | 172375 | 140857 | 142682 | 386739 | 391523 |
| Number of GIAB-only calls in GIAB highconf bed | 90722 | 122544 | 121076 | 121207 | 124715 | 195277 | 163467 | 157494 | 111787 |
| Number of concordant calls that are filtered by the GIAB highconf bed | 12 | 0 | 736918 | 657715 | 608137 | 53460 | 51255 | 48696 | 45779 |

### High Concordance with Illumina Platinum Genomes

The bottom section of Table 1 also shows the increasing number of concordant and decreasing number of discordant calls over multiple versions of our HG001 callset compared to the Illumina Platinum Genomes 2016-v1.0 callset (PG).^6^ PG is a valuable benchmark for our calls because it uses a phased pedigree analysis to arbitrate between a different set of input variant calls to arrive at high-confidence vcf and bed files. This provides orthogonal confirmation of our calls, since the PG phased pedigree analysis may identify different biases from our method. PG contains a larger number of high-confidence variant calls, even relative to our newest v3.3.2 calls for HG001. However, when comparing v3.3.2 to PG within both bed files, we found that the majority of differences were places where v3.3.2 was correct and PG had a partial complex or compound heterozygous call (e.g., Fig. 2), or where the PG bed file partially overlapped with a true deletion that v3.3.2 called correctly. To reduce the number of these problematic sites, we contracted each of the PG high-confidence regions by 50bp on each side, which is similar to how we remove 50bp on each side of uncertain variants (e.g., locus (1) in Fig. 1b). Because PG has many more uncertain variants, contracting the PG high-confidence regions by 50bp reduces the number of variants in the PG bed by 32%, but it also eliminates 93% of the differences between v3.3.2 and PG in GRCh37, so that 61 PG-only and 47 v3.3.2-only calls remain in both bed files.

**Fig. 2:**
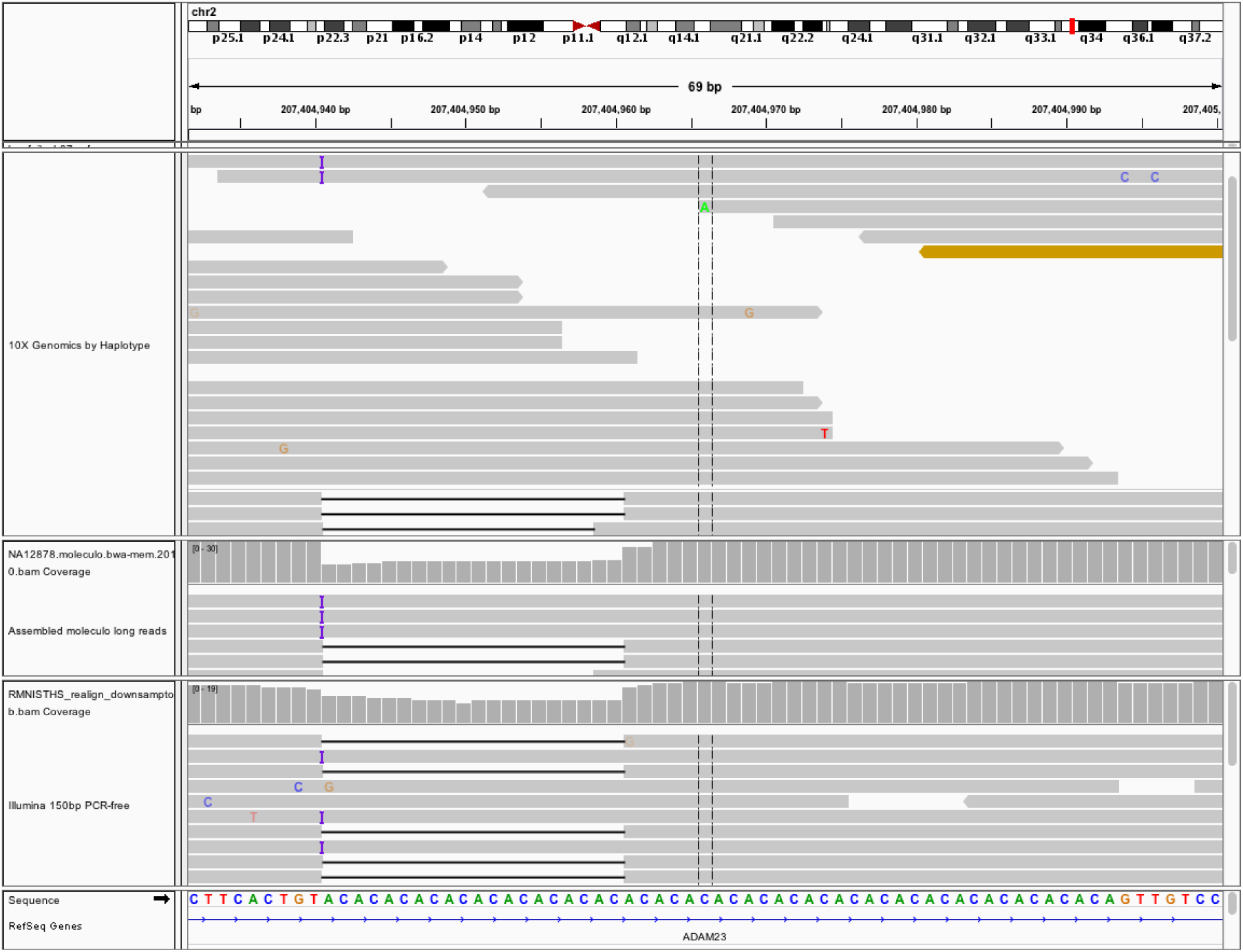
Compound heterozygous insertion and deletion in HG001 in a tandem repeat at 2:207404940 (GRCh37), for which Illumina Platinum Genomes only calls a heterozygous deletion. When a callset with the true compound heterozygous variant is compared to Platinum Genomes, it is counted as both a FP and a FN. Both the insertion and deletion are supported by PCR-free Illumina (bottom) and Moleculo assembled long reads (middle), and reads assigned haplotype 1 in 10x support the insertion and reads assigned haplotype 2 in 10x support the deletion (top).

After manually curating the remaining 108 differences between v3.3.2 and PG in GRCh37, v3.3.2 has 5 clear FNs, 1 clear FP, 2 unclear potential FNs, and 6 unclear potential FPs in the NIST high-confidence regions (see Supplementary Table 1 and supplementary text for details). Based on 3529641 variants concordant between NIST and PG, there would be about 2 FPs and 2 FNs per million true variants. Since these are only in the PG high-confidence regions, these counts are very likely lower than the overall counts of errors in v3.3.2, which are difficult to estimate. In addition, our estimates of error rates may be biased because we fixed problems in previous versions of the integration based on errors we found upon manually inspecting differences between our calls and PG (primarily in chromosomes 1 and 20). For these reasons, the true error rate of our calls is likely to be higher than 2 FPs and 2 FNs per million true variants.

### Trio Mendelian inheritance analysis

For the Ashkenazi trio, we have separately developed high-confidence calls and regions for the son (HG002), father (HG003), and mother (HG004), which allows us both to phase some variants in the son (see below) and to determine variants that have a genotype pattern inconsistent with that expected by Mendelian inheritance. Because variants are called independently in the three individuals, complex variants can be represented differently in the 3 individuals; the majority of apparent Mendelian violations found by naive methods are actually just different representations of complex variants in different individuals. We used a new method based on rtg-tools vcfeval to harmonize representation of variants across the 3 individuals, similar to a recent publication.^14^ For our GRCh37 high-confidence calls, this decreased the number of apparent Mendelian violations inside all 3 high-confidence regions from 30151/4390900 (0.68%) to 2038/4383371 (0.05%). Of the 2038 violations, 1323 (1110 SNPs and 213 indels) were likely to be either cell-line somatic or germline *de novo* mutations, because the son was heterozygous and the parents were homozygous reference. The number of SNPs is about 10% higher than the 1001 de novo SNPs found by 1000 Genomes for NA12878/HG001, and higher than the 678 de novo SNPs they found for NA19240.^15^ Note that our data was much higher coverage than the 1000 Genomes data, but we also excluded any variants below 0.2 allele fraction from our high-confidence regions, so variants in a small fraction of the cells, as well as other variants outside our high-confidence regions in any member of the trio, are not included in our counts.

Upon manual inspection of 10 random de novo SNPs (see Supplementary Table 1 for details), all were clearly present in multiple technologies for the son and not in any data for the parents, and they were in all or part of only one 10x haplotype. At one randomly inspected site, there unexpectedly appear to be two different strongly supported de novo SNPs on the same haplotype at 15:83717817, and both SNPs are supported by all technologies. Upon manual inspection of 10 random de novo indels, 7 appeared to be true de novo indels (most in homopolymers or tandem repeats), 1 was a correctly called de novo SNP phased with an indel inherited from HG004, and 2 were indels that were correctly called in HG002 but missed in one of the parents due to an issue in converting gvcf to vcf that excludes low QUAL variants even with high GQ. The remaining 715 non-de novo Mendelian violations were 91% indels. They mostly resulted from errors in the integration process in one of the individuals, often related to incorrectly calling two different alleles in the one of the individuals in the same repetitive region. We exclude 50bp on either side of these 715 violations from our final high-confidence bed files for all 3 individuals.

Based on the Mendelian analysis, there may be 16 SNP and 150 indel errors per million variants in one of the 3 individuals before excluding these sites. Note that we exclude the errors found by the Mendelian analysis from our final high-confidence bed files, but it is likely that the error modality that caused these Mendelian errors also caused other errors that were not found to be Mendelian inconsistent. Many other error modalities can be Mendelian consistent (e.g., systematic errors that cause everyone to be heterozygous and systematic errors that cause the wrong allele to be called), so these error estimates should be interpreted with caution.

### Performance metrics can vary significantly when comparing to different benchmark callsets

When comparing a method’s callset against imperfect and incomplete benchmark callsets, it is important to recognize that estimates of precision and recall/sensitivity may differ from the true precision and recall for the method for all regions of the genome and all types of variants. If there are insufficient examples of variants of particular types, estimates of precision and recall will have high uncertainties. In addition, estimates of precision and recall can be biased (1) by errors in the benchmark callset, (2) by using comparison methods that do not compare differing representations of complex variants, and (3) by biases in the benchmark towards easier variants and genome contexts. We demonstrate that these biases are particularly apparent for some challenging variant types and genome contexts by benchmarking a standard bwa-GATK callset against our new calls, calls from our 2014 publication, and Platinum Genomes.

We examine 3 striking examples in coding regions, complex variants, and decoy-associated regions (see Supplementary Table 1 and supplementary text for details): (1) When bwa-GATK is compared to PG the sensitivity to SNPs in RefSeq coding regions is 97.98%, whereas when bwa-GATK is compared to v3.3.2 the sensitivity to SNPs in RefSeq coding regions is 99.96% (a FN rate estimate 62 times lower). Upon manual curation of 10 putative FNs from the PG comparison, all were in difficult-to-map regions, 8 had some long read support, and 2 appeared to be inaccurate PG calls. (2) Similarly, we manually inspected a random subset of putative FP indels that were in or near compound heterozygous variants. When compared to PG, the precision for these was 93%, whereas when compared to v3.3.2 the precision was 95%.

When inspecting 10 random GATK FPs compared to v3.3.2, we found 9 were likely errors in bwa-GATK mis-genotyping the true compound heterozygous event, and 1 was a likely mis-genotype in v3.3.2. When inspecting 10 random GATK FPs compared to PG, we found 4 were likely errors in bwa-GATK mis-genotyping the true compound heterozygous event, and 6 were accurate compound heterozygous events that were only partially called or missed by PG. In this case, the v3.3.2 appears more useful than PG at finding real FPs, because many of the putative FPs vs PG are not actually errors. (3) Finally, we examined regions homologous to the hs37d5 decoy sequence from 1000 Genomes. We compared recall, precision, and fraction not assessed for the decoy-associated regions when comparing bwa-GATK calls that did not include the decoy when mapping (bwa-GATK-nodecoy) to our v3.3.2, v2.18, and PG high-confidence calls. The precision for bwa-GATK-nodecoy is 67% vs. v3.3.2, whereas it is 91% vs. v2.18 and 93% vs. PG. The much higher precision vs v2.18 or PG is a result of many of the decoy-associated variants being excluded from the high-confidence regions of v2.18 and PG. Assessing these variants is important because they make up 7123 of the 16452 FPs (43%) vs. v3.3.2 for this callset, even though variants in decoy-associated regions only make up 0.5% of TPs. However, in all of these cases, the true precision and recall of bwa-GATK is unknown outside the v3.3.2 and PG bed files.

## Discussion

Here, we present new, robust methods to form benchmark high-confidence SNP, indel, and homozygous reference calls from multiple sequencing datasets from the same individual. By developing reproducible integration methods that work on arbitrary genomes with multiple datasets, we were able to form high-confidence benchmark calls for both GRCh37 and GRCh38 for four additional individuals that are new whole genome Reference Materials from NIST. Not only are the integration methods sufficiently robust to form calls on multiple genomes, but we also show that these calls are more accurate and more comprehensive than our previous high-confidence callsets when benchmarked against other high-quality callsets. We show that when using our high-confidence calls with best practices for benchmarking established by the Global Alliance for Genomics and Health,^8^ the majority of putative FPs and FNs are errors in the callset being benchmarked. Therefore, they can be reliably used to identify errors in a sequencing and bioinformatics process.

One important limitation of high-confidence callsets like ours is that they will tend to exclude more difficult types of variation and regions of the genome. When new methods that attempt to call difficult variants are compared to our current callsets, they may have lower accuracy in our high-confidence regions relative to more mature methods, even though they may have much higher accuracy in difficult regions relative to mature methods. For example, as graph-based variant calling methods are developed, they may be able to make much better calls in regions with many ALT haplotypes like the Major Histocompatibility Complex (MHC) even if they have lower accuracy within our high-confidence regions. Therefore, it can be important to consider variant calls outside our high-confidence regions when evaluating the accuracy of different variant calling methods.

To assure that our high-confidence calls continue to be useful as sequencing and analysis methods improve, we are exploring several approaches for characterizing more difficult regions of the genome. For HG001, pedigree information can be used to filter out likely errors in difficult regions, particularly in homopolymers, and phased pedigree-based calls from Illumina Platinum Genomes and Real Time Genomics could be used to supplement our high-confidence calls. Long reads and linked reads show promise in enabling high-confidence calls in difficult-to-map regions of the genome. In addition, some clinical laboratories have designed assays to target difficult-to-sequence genes, and their data could be incorporated into our high-confidence calls. The methods presented here are a framework to integrate all of these data types, so long as variant calls with high-sensitivity in a specified set of genomic regions could be generated, and filters could be applied to give high specificity.

The Genome in a Bottle Consortium is planning to select additional genomes for deep characterization, because the current benchmark genomes represent limited ancestry groups. The Consortium is also evaluating the possibility of adding at least one large family to enable segregation consistency validation of variants. Future work is still needed to understand whether variant calling performance varies between ancestry groups.

Finally, to ensure these data are an enduring resource, we have made available an online form, where we and others can enter potentially questionable sites in our high-confidence regions (https://goo.gl/forms/zvxjRsYTdrkhqdzM2). The results of this form are public and updated in real-time so that anyone can see where others have manually reviewed or interrogated the evidence at any site (https://docs.google.com/spreadsheets/d/1kHgRLinYcnxX3-ulvijf2HrIdrQWz5R5PtxZS-_s6ZM/edit?usp=sharing).

## Acknowledgments

We thank the many contributors to Genome in a Bottle Consortium discussions. We especially thank Rafael Saldana and the Sentieon team for advice on running the Sentieon pipeline, Andrew Carroll and the DNAnexus team for advice on implementing the pipeline in DNAnexus, Fiona Hyland, Srinka Ghosh, Keyan Zhao, and John Bodeau at ThermoFisher for advice integrating Ion exome and SOLiD genome data, Deanna Church and Valerie Schneider for helpful discussions about GRCh38, and many individuals for providing feedback about the current version and previous versions of our calls. Certain commercial equipment, instruments, or materials are identified to specify adequately experimental conditions or reported results. Such identification does not imply recommendation or endorsement by the National Institute of Standards, nor does it imply that the equipment, instruments, or materials identified are necessarily the best available for the purpose.

## Data availability

The high-confidence vcf and bed files from this manuscript are available in the NISTv3.3.2 directory under each genome on the GIAB FTP release folder ftp://ftp-trace.ncbi.nlm.nih.gov/giab/ftp/release/, and in the future updated calls will be in the “recent” directory under each genome. The data used in this manuscript and other datasets for these genomes are available in ftp://ftp-trace.ncbi.nlm.nih.gov/giab/ftp/data/, and the raw data used here and additional data for these genomes are also available in the NCBI BioProject PRJNA200694.

## Author contributions

JMZ, LT, and MS wrote the manuscript.

JMZ, JM, FMD, MS, and HP designed and implemented the integration process.

HH, JM, and JMZ analyzed and integrated the 10x Genomics data.

RT, JM, and JMZ analyzed and integrated the Complete Genomics data.

SAI, LT, FMD, JM, and JMZ designed and implemented the phasing and robust trio analysis.

CYM, JM, and JMZ designed and implemented the robust GRCh38 liftover analysis.

CX and SS managed and analyzed data.

All authors contributed to GIAB discussions planning this work.

## Online Methods

### Sequencing Datasets

In contrast to our previous integration process,^1^ which used sequencing data that was generated from DNA from various growths of the Coriell cell line GM12878, most of the sequencing data used in this work were generated from NIST RMs, which are DNA extracted from a single large batch of cells that was mixed prior to aliquoting.^10^ These datasets, except for the HG001 SOLiD datasets, are described in detail in a Genome in a Bottle data publication.^10^

For these genomes, we used datasets from several technologies and library preparations:

1. ∼300x paired end whole genome sequencing with 2×148bp reads with ∼550bp insert size from the HiSeq 2500 in Rapid Mode with v1 sequencing chemistry (HG001 and AJ Trio).
2. ∼300x paired end whole genome sequencing with 2×250bp reads with ∼550bp insert size from the HiSeq 2500 in Rapid Mode with v2 sequencing chemistry (Chinese son).
3. ∼45x paired end whole genome sequencing with 2×250bp reads with ∼400bp insert size from the HiSeq 2500 in Rapid Mode with v2 sequencing chemistry (AJ Trio).
4. ∼15x mate-pair whole genome sequencing with 2×100bp reads with ∼6000bp insert size from the HiSeq 2500 in High Throughput Mode with v2 sequencing chemistry (AJ Trio and Chinese son).
5. ∼100x paired end 2×29bp whole genome sequencing from Complete Genomics v2 chemistry (all genomes)
6. ∼1000x single end exome sequencing from Ion PI(tm) Sequencing 200 Kit v4 (all genomes)
7. Whole genome sequencing from SOLiD 5500W. For HG001, two whole genome sequencing datasets were generated: ∼12x 2×50bp paired end and ∼12x 75bp single end with error-correction chemistry (ECC). For the AJ son and Chinese son, ∼42x 75bp single end sequencing (without ECC) was generated.
8. 10x Genomics Chromium whole genome sequencing (∼25x per haplotype from HG001 and AJ son, and ∼9x per haplotype from AJ parents). These are the only data not from the NIST RM batch of DNA because longer DNA from the cell lines resulted in better libraries

### Implementation of analyses and source code

Most analyses were performed using apps or applets on the DNAnexus data analysis platform (DNAnexus, Inc., Mountain View, CA), except for mapping of all datasets and variant calling for Complete Genomics and Ion exome, since these steps were performed previously. The apps and applets used in this work are included in GitHub (https://github.com/jzook/genome-data-integration/tree/master/NISTv3.3.2). They run on an Ubuntu 12.04 machine on Amazon Web Services EC2. The apps and applets are structured as:

1. dxapp.json specifies the input files and options, output files, and any dependencies that can be installed via the Ubuntu command *apt*.
2. src/code.sh contains the commands that are run
3. resources/contains compiled binary files, scripts, and other files that are used in the applet

The commands were run per chromosome in parallel using the DNAnexus command line interface, and these commands are listed at https://github.com/jzook/genome-data-integration/tree/master/NISTv3.3.2/DNAnexusCommands/batch_processing_commands.

### Illumina analyses

The Illumina fastq files were mapped using novoalign -d <reference.ndx> -f <read1.fastq.gz> <read2.fastq.gz> -F STDFQ --Q2Off -t 400 -o SAM -c 10 (Novocraft Technologies, Selangor, Malaysia).

and resulting BAM files were created, sorted, and indexed with samtools version 0.1.18. Bam files are available under:

ftp://ftp-trace.ncbi.nlm.nih.gov/giab/ftp/data/NA12878/NIST_NA12878_HG001_HiSeq_300x/

ftp://ftp-trace.ncbi.nlm.nih.gov/giab/ftp/data/AshkenazimTrio/HG002_NA24385_son/NIST_HiSeq_HG002_Homogeneity-10953946/

ftp://ftp-trace.ncbi.nlm.nih.gov/giab/ftp/data/AshkenazimTrio/HG002_NA24385_son/NIST_Illumina_2×250bps/

ftp://ftp-trace.ncbi.nlm.nih.gov/giab/ftp/data/AshkenazimTrio/HG003_NA24149_father/NIST_HiSeqHG003_Homogeneity-12389378/

ftp://ftp-trace.ncbi.nlm.nih.gov/giab/ftp/data/AshkenazimTrio/HG003_NA24149_father/NIST_Illumina_2×250bps/

ftp://ftp-trace.ncbi.nlm.nih.gov/giab/ftp/data/AshkenazimTrio/HG004_NA24143_mother/NIST_HiSeqHG004_Homogeneity-14572558/

ftp://ftp-trace.ncbi.nlm.nih.gov/giab/ftp/data/AshkenazimTrio/HG004_NA24143_mother/NIST_Illumina_2×250bps/

ftp://ftp-trace.ncbi.nlm.nih.gov/giab/ftp/data/ChineseTrio/HG005_NA24631_son/HG005_NA24631_son_HiSeq_300x

Variants were called using both GATK HaplotypeCaller v3.5 ^16,17^ and Freebayes 0.9.20 ^18^ with high-sensitivity settings. Specifically, for HaplotypeCaller, special options were “-stand_call_conf 2 -stand_emit_conf 2 -A BaseQualityRankSumTest -A ClippingRankSumTest -A Coverage -A FisherStrand -A LowMQ -A RMSMappingQuality -A ReadPosRankSumTest -A StrandOddsRatio - A HomopolymerRun -A TandemRepeatAnnotator”. The gVCF output was converted to VCF using GATK Genotype gVCFs for each sample independently. For freebayes, special options were “-F-m 0 --genotype-qualities”.

For freebayes calls, GATK CallableLoci v3.5 was used to generate callable regions with at least 20 reads with mapping quality of at least 20 (to exclude regions where heterozygous variants may be missed), and with coverage less than two times the median (to exclude regions likely to be duplicated or have mis-mapped reads). Because parallelization of freebayes infrequently causes conflicting variant calls at the same position, 2 variants at the same position are removed from the callable regions.

For GATK calls, the gVCF output from GATK was used to define the callable regions. In general, reference regions and variant calls with GQ<60 were excluded from the callable regions, excluding 50bp on either side of low GQ reference regions and low GQ variant calls. Excluding 50bp minimizes artifacts introduced when integrating partial complex variant calls. The exception to this rule is that reference assertions with GQ<60 are ignored if they are within 10bp of start or end an indel with GQ>60, because upon manual inspection GATK often calls some reference bases with a low GQ near true indels even when the reference bases should have high GQ, and excluding 50bp regions around these bases excluded many true high-confidence indels. The gVCF from GATK is used rather than CallableLoci because it provides a sophisticated interrogation of homopolymers and tandem repeats and excludes regions if insufficient reads completely cross the repeats.

### Complete Genomics analyses

Complete Genomics data was mapped and variants were called using v2.5.0.33 of the standard Complete Genomics pipeline.^19^ Only the vcfBeta file was used in the integration process, because it contains both called variants and “no call” regions similar to gVCF. vcfBeta files are available under:

ftp://ftp-trace.ncbi.nlm.nih.gov/giab/ftp/data/NA12878/analysis/CompleteGenomics_RMDNA_11272014/

ftp://ftptrace.ncbi.nlm.nih.gov/giab/ftp/data/AshkenazimTrio/analysis/CompleteGenomics_RefMaterial_SmallVariants_CGAtools_08082014/

ftp://ftp-trace.ncbi.nlm.nih.gov/giab/ftp/data/ChineseTrio/analysis/CompleteGenomics_HanTrio_RMDNA_08082014/

A python script from Complete Genomics (vcf2bed.py) was used to generate callable regions, which exclude regions with no calls or partial no calls in the vcfBeta. To minimize integration artifacts around partial complex variant calls, 50bp padding was subtracted from both sides of callable regions using bedtools slopBed. In addition, the vcfBeta file was modified to remove unnecessary lines and fill in the FILTER field using a python script from Complete Genomics (vcfBeta_to_VCF_simple.py). This process was performed on DNAnexus; an example command for chr20 is “dx run GIAB:/Workflow/integration-prepare-cg-ivcf_in=/NA12878/Complete_Genomics/vcfBeta-GS000025639-ASM.vcf.bz2-ichrom=20--destination=/NA12878/Complete_Genomics/”

### Ion exome analyses

The Ion exome data BaseCalling and alignment were performed on a Torrent Suite v4.2 server (ThermoFisher Scientific). Variant calling was performed using Torrent Variant Caller v4.4, and the TSVC_variants_defaultlowsetting.vcf was used as a sensitive variant call file.

GATK CallableLoci v3.5 was used to generate callable regions with at least 20 reads with mapping quality of at least 20.

These callable regions were intersected with the targeted regions bed file for the Ion Ampliseq exome assay available at ftp://ftp-trace.ncbi.nlm.nih.gov/giab/ftp/data/AshkenazimTrio/analysis/IonTorrent_TVC_03162015/AmpliseqExome.20141120_effective_regions.bed. In addition, 50bp on either side of compound heterozygous sites were removed from the callable regions, and these sites were removed from the vcf to avoid artifacts around homopolymers. This process was performed on DNAnexus, with the command for chr20:

“dx run GIAB:/Workflow/integration-prepare-ion - ivcf_in=/NA12878/Ion_Torrent/TSVC_variants_defaultlowsetting.vcf - icallablelocibed=/NA12878/Ion_Torrent/callableLoci_output/HG001_20_hs37d5_IonExome_callab leloci.bed -itargetsbed=/NA12878/Ion_Torrent/AmpliseqExome.20141120_effective_regions.bed - ichrom=20 --destination=/NA12878/Ion_Torrent/Integration_prepare_ion_output/”

### SOLiD analyses

The SOLiD xsq files were mapped with LifeScope v2.5.1 (ThermoFisher Scientific). Bam files are available under:

ftp://ftp-trace.ncbi.nlm.nih.gov/giab/ftp/data/NA12878/NIST_NA12878_HG001_SOLID5500W/

ftp://ftp-trace.ncbi.nlm.nih.gov/giab/ftp/data/AshkenazimTrio/HG002_NA24385_son/NIST_SOLiD5500W

ftp://ftp-trace.ncbi.nlm.nih.gov/giab/ftp/data/ChineseTrio/HG005_NA24631_son/NIST_SOLiD5500W/

Variants were called using GATK HaplotypeCaller v3.5 with high-sensitivity settings. Specifically, for HaplotypeCaller, special options were “-stand_call_conf 2 -stand_emit_conf 2 -A BaseQualityRankSumTest -A ClippingRankSumTest -A Coverage -A FisherStrand -A LowMQ -A RMSMappingQuality -A ReadPosRankSumTest -A StrandOddsRatio -A HomopolymerRun -A TandemRepeatAnnotator”. The gVCF output was converted to VCF using GATK Genotype gVCFs for each sample independently.

For SOLiD, all regions were considered “not callable” because biases are not sufficiently well understood, so SOLiD only provided support from an additional technology when finding training sites and when annotating the high-confidence vcf.

### 10x Genomics analyses

The 10x Genomics Chromium fastq files were mapped and reads were phased using LongRanger v2.1 (10x Genomics, Pleasanton, CA). As a new technology, variant calls are integrated in a conservative manner, requiring clear support for a homozygous variant or reference call from reads on each haplotype. Bamtools was used to split the bam file into two separate bam files with reads from each haplotype (HP tag values 1 and 2), ignoring reads that were not phased. GATK HaplotypeCaller v3.5 was used to generate gvcf files from each haplotype separately. A custom perl script was used to parse the gvcf files, excluding regions with DP<6, DP > 2x median coverage, heterozygous calls on a haplotype, or homozygous reference or variant calls where the likelihood was <99 for homozygous variant or reference, respectively. The union of these regions from both haplotypes plus 50bp on either side was excluded from the callable regions.

### Excluding challenging regions for each callset

As an enhancement in our new integration methods (v3.3+), we now exclude difficult regions per callset, rather than at the end for all callsets. We used prior knowledge as well as manual inspection of differences between callsets to determine which regions should be excluded from each callset. For example, we exclude tandem repeats and perfect or imperfect homopolymers >10bp for callsets with reads <100bp or from technologies that use PCR (all methods except Illumina PCR-free). We exclude segmental duplications and regions homologous to the decoy hs37d5 or ALT loci from all short read methods (all methods except 10x Genomics linked reads). The regions excluded from each callset are specified in the “CallsetTables” input into the integration process, where a 1 in a column indicates the bed file regions are excluded from that callset’s callable regions, and a 0 indicates the bed file regions are not excluded for that callset. Callset tables are available at https://github.com/jzook/genome-data-integration/tree/master/NISTv3.3.2/CallsetTables.

### Integration process to form high-confidence calls and regions

Our integration process is summarized in Fig. 1 and detailed in the outline in the Supplementary Methods. Similar to our previous work for v2.18/v2.19,^1^ the first step in our integration process is to use preliminary “concordant” calls to train a machine learning model that finds outliers. Callsets with a genotype call that has outlier annotations are not trusted. For training variants, we use genotype calls that are supported by at least 2 technologies, not including sites if another dataset contradicts the call or is missing the call and the call is within that callset’s callable regions. To do this, we normalize variants using vcflib vcfallelicprimitives and then generate a union vcf using vcflib vcfcombine. The union vcf contains separate columns from each callset, and we annotate the union vcf with vcflib vcfannotate to indicate whether each call falls outside the callable bed file from each dataset.

To find outliers, we selected annotations from the vcf INFO and FORMAT fields and which tail of the distribution that we expected to be associated with bias for each callset (files describing annotations for each caller are at https://github.com/jzook/genome-data-integration/tree/master/NISTv3.3.2/AnnotationFiles). Then, we used a simple one-class model that treats each annotation independently and finds the cut-off (for single tail) or cut-offs (for both tails) outside which (5/*a*) % of the training calls lie, where *a* is the number of annotations for each callset. For each callset, we find sites that fall outside the cut-offs or that are already filtered, and we generate a bed file that contains the call with 50bp added to either side to account for different representations of complex variants. We again annotate the union vcf with this “filter bed file” from each callset, which we next use in addition to the “callable regions” annotations (Fig. 1b).

To generate the high-confidence calls, we run the same integration script with the new union vcf annotated with both “callable regions” from each dataset and “filtered regions” from each callset (Fig. 1c). The integration script outputs high-confidence calls that meet all of the following criteria (an appropriate output vcf FILTER status is given for sites that don’t meet each criterion):

1. Genotypes agree between all callsets for which the call is callable and not filtered. This includes “implied homozygous reference calls”, where a site is missing from a callset and it is within that callset’s callable regions and no filtered variants are within 50bp (otherwise, FILTER status “discordantunfiltered”, e.g., example (2) in Fig. 1c).
2. The call from at least one callset is callable and not filtered (otherwise, FILTER status “allfilteredbutagree” or “allfilteredanddisagree”, e.g., example (4) in Fig. 1c).
3. The sum of genotype qualities for all datasets supporting this genotype call is >=70. This sum includes only the first callset from each dataset and includes all datasets supporting the call even if they are filtered or outside the callable bed (otherwise, FILTER status “GQlessthan70”).
4. If the call is homozygous reference, then no filtered calls at the location can be indels >9bp. Implied homozygous reference calls (i.e., there is no variant call and it is in the callset’s callable regions are sometimes unreliable for larger indels, because callsets will sometimes miss the evidence for large indels and not call a variant (otherwise, FILTER status “questionableindel”).
5. The site is not called only by Complete Genomics and completely missing from other callsets, since these sites tended to be systematic errors in repetitive regions upon manual curation (otherwise, FILTER status “cgonly”).
6. For sites where the high-confidence call is heterozygous, none of the filtered calls are homozygous variant, since these are sometimes genotyping errors (otherwise, FILTER status “discordanthet”).
7. Heterozygous calls where the net allele balance across all unfiltered datasets is <0.2 or >0.8 when summing support for REF and ALT alleles (otherwise, FILTER status “alleleimbalance”).

To calculate the high-confidence regions bed file, the following steps are performed:

1. Find all regions that are covered by at least one dataset’s callable regions bed file.
2. Subtract 50bp on either side of all sites that could not be determined with high confidence.

We chose to use a simple, one-class model in this work in place of the more sophisticated Gaussian Mixture Model (GMM) used in v2.19 to make the integration process more robust and reproducible. In we way we used the GMM in v2.18 and v2.19, GATK Variant Quality Score Recalibration would sometimes not find a fit to the data, so that parameters would need to be manually modified, making the integration process less robust and reproducible. At times the GMM also appeared to overfit the data, filtering sites for unclear reasons. The GMM also was designed for GATK, ideally for vcf files containing many individuals, unlike our single sample vcf files. Our one-class model used in v3.3.2 was more easily adapted to filter on annotations in the FORMAT and INFO fields in vcf files produced by the variety of variant callers and technologies used for each sample. Future work could be directed towards developing more sophisticated models that learn which annotations are more useful, how to define the callable regions, and how annotations may have different values for different types of variants and sizes.

The integration process described in this section is implemented in the applet nist-integration-v3.3.2-anyref (https://github.com/jzook/genome-data-integration/tree/master/NISTv3.3.2/DNAnexusApplets/nist-integration-v3.3.2-anyref). The one-class filtering model to find outliers is a custom perl script (nist-integration-v3.3.2-anyref/resources/usr/bin/VcfOneClassFiltering_v3.3.pl). The integration and arbitration processes to (1) find genotype calls supported by two technologies for the training set and (2) find high-confidence genotype calls are both implemented in a perl script (nist-integration-v3.3.2-anyref/resources/usr/bin/VcfClassifyUsingFilters_v3.3.pl).

### Candidate structural variants to exclude from high-confidence regions

For HG001, we had previously used all HG001 variants from dbVar, which conservatively excluded about 10% of the genome, because dbVar contained any variants anyone ever submitted for HG001. Since then, several SV callsets from different technologies have been generated for HG001, and we now use these callsets instead of dbVar. Specifically, we include union of all calls <1Mbp in size from:

1. The union of several PacBio SV calling methods, including filtered sites (ftp://ftp-trace.ncbi.nlm.nih.gov/giab/ftp/data/NA12878/NA12878_PacBio_MtSinai/NA12878.sorted.vcf.gz).
2. The PASSing calls from MetaSV, which looks for support from multiple types of Illumina calling methods (ftp://ftp-trace.ncbi.nlm.nih.gov/giab/ftp/technical/svclassify_Manuscript/Supplementary_Information/metasv_trio_validation/NA12878_svs.vcf.gz).
3. Calls with support from multiple technologies from svclassify (ftp://ftp-trace.ncbi.nlm.nih.gov/giab/ftp/technical/svclassify_Manuscript/Supplementary_Information/Personalis_1000_Genomes_deduplicated_deletions.bed and ftp://ftp-trace.ncbi.nlm.nih.gov/giab/ftp/technical/svclassify_Manuscript/Supplementary_Information/Spiral_Genetics_insertions.bed).

For the AJ trio, we use the union of calls from 11 callers using 5 technologies. Specifically, we include all calls >50bp and <1Mbp in size from:

1. PacBio callers: sniffles, Parliament, CSHL assembly, SMRT-SV dip, and MultibreakSV
2. Illumina callers: Cortex, Spiral Genetics, Jitterbug, and Parliament
3. BioNano haplo-aware calls
4. 10x GemCode
5. Complete Genomics highConfidenceSvEventsBeta and mobileElementInsertionsBeta

For deletions, we add 100bp to either side of the called region, and for all other calls (mostly insertions), we add 1000bp to either side of the called region. This padding helps to account for imprecision in the called breakpoint, complex variation around the breakpoints, and potential errors in large tandem duplications that are reported as insertions.

### GRCh38 integration

To develop high-confidence calls for GRCh38, we use similar methods to GRCh37 except for data that was not mapped to GRCh38. We are able to map reads and call variants directly on GRCh38 for Illumina and 10x, but native GRCh38 pipelines were not available for Complete Genomics, Ion exome, and SOLiD data. For Illumina and 10x data, variant calls were made similarly to GRCh37 but from reads mapped to GRCh38 with decoy but no alts (ftp://ftp.ncbi.nlm.nih.gov/genomes/all/GCA/000/001/405/GCA_000001405.15_GRCh38/seqs_for_alignment_pipelines.ucsc_ids/GCA_000001405.15_GRCh38_no_alt_plus_hs38d1_analysis_set.fna.gz). For Complete Genomics, Ion exome, and SOLiD data, vcf and callable bed files were converted from GRCh37 to GRCh38 using the tool GenomeWarp (https://github.com/verilylifesciences/genomewarp). This tool converts vcf and callable bed files in a conservative and sophisticated manner, accounting for base changes that were made between the two references. Modeled centromere (genomic_regions_definitions_modeledcentromere.bed) and heterochromatin (genomic_regions_definitions_heterochrom.bed) regions are explicitly excluded from the high-confidence bed (available under ftp://ftp-trace.ncbi.nlm.nih.gov/giab/ftp/release/NA12878_HG001/NISTv3.3.2/GRCh38/supplementaryFiles/).

For HG001 Illumina and 10x data, variants were called using GATK HaplotypeCaller v3.5 similarly to all GRCh37 genomes, but for the other genomes’ Illumina and 10x datasets, the Sentieon haplotyper was used instead. (Sentieon, Inc., Mountain View, CA). This tool was designed to give the same results more efficiently than GATK HaplotypeCaller v3.5, except that it does not downsample reads in high coverage regions, so that the resulting variant calls are deterministic.

### Trio Mendelian analysis and phasing

We performed a Mendelian analysis and phasing for the AJ Trio using rtg-tools v3.7.1. First, we harmonize the representation of variants across the trio using an experimental mode of rtg-tools vcfeval, so that different representations of complex variants do not cause apparent Mendelian violations. After merging the 3 individuals’ harmonized vcfs into a multi-sample vcf, we change missing genotypes for individuals to homozygous reference, since we later subset by high-confidence regions (missing calls are implied to be homozygous reference in our high-confidence regions). Then, we use rtg-tools mendelian to phase variants in the son and find Mendelian violations. Finally, we subset the Mendelian violations by the intersection of the high-confidence bed files from all three individuals. We also generate new high-confidence bed files for each individual that exclude 50bp on each side of Mendelian violations that are not apparent de novo mutations. This analysis and the phase transfer below is performed in the DNAnexus applet *trio-harmonize-mendelian* (https://github.com/jzook/genome-data-integration/tree/master/NISTv3.3.2/DNAnexusApplets/trio-harmonize-mendelian).

### Phase transfer

Our integration methods only supply local phasing where GATK HaplotypeCaller (or Sentieon haplotyper) is able to phase the variants. For HG001/NA12878 and the AJ son (HG002), pedigree-based phasing information can supply global phasing of maternal and paternal haplotypes.

For HG001, Real Time Genomics (ftp://ftp-trace.ncbi.nlm.nih.gov/giab/ftp/data/NA12878/analysis/RTG_Illumina_Segregation_Phasing_05122016/) and Illumina Platinum Genomes (ftp://ftp-trace.ncbi.nlm.nih.gov/giab/ftp/technical/platinum_genomes/2016-1.0/) have generated phased variant calls by analyzing her 17-member pedigree. The Platinum Genomes calls are phased in the order paternal|maternal, and the RTG phasing is not ordered, so the phasing from Platinum Genomes was added first. First, we archive existing local phasing information to the IGT and IPS fields. Then, we use the rtg-tools vcfeval software phase-transfer mode to take the phasing first from Platinum Genomes and add it to the GT field of our HG001 high-confidence vcf. We then use the rtg-tools vcfeval’s phase-transfer mode to take the phasing from RTG and add it to variants that were not phased by Platinum Genomes, flipping the phasing to the order paternal|maternal where necessary. Variants with phasing from PG or RTG are given a PS=PATMAT. Variants that were not phased by either PG or RTG but are homozygous variant are given GT=1|1 and PS=HOMVAR. Variants that were not phased by either PG or RTG but were phased locally by GATK HaplotypeCaller or Sentieon haplotyper are given the phased GT and PS from the variant caller. This phase transfer is performed in 2 steps with the applet *phase-transfer*.

For HG002, we apply the phasing from the trio analysis using rtg-tools vcfeval phase-transfer like above. Variants with trio-based phasing are given a PS=PATMAT. Variants that were not phased by the trio but are homozygous variant are given GT=1|1 and PS=HOMVAR. Variants that were not phased by the trio but were phased locally by GATK HaplotypeCaller or Sentieon haplotyper are given the phased GT and PS from the variant caller. This phase transfer is performed in the applet *trio-harmonize-mendelian*.

### Comparisons to other callsets

The Global Alliance for Genomics and Health formed a Benchmarking Team (https://github.com/ga4gh/benchmarking-tools) to standardize performance metrics and develop tools to compare different representations of complex variants. We have used one of these tools, vcfeval (https://github.com/RealTimeGenomics/rtg-tools) from rtg-tools-3.6.2 with --ref-overlap, to compare our high-confidence calls to other vcfs. After performing the comparison, we subset the true positives, false positives, and false negatives by our high-confidence bed file and then by the bed file accompanying the other vcf (if it has one). We then manually inspect alignments from a subset of the putative false positives and false negative and record whether we determine our high-confidence call is likely correct, if we understand why the other callset is incorrect, if the evidence is unclear, if it is in a homopolymer, and other notes.

For HG001, we compare to these callsets:

1. NISTv2.18 high-confidence calls and bed file we published previously
2. Platinum Genomes 2016-1.0, with a modified high-confidence bed file that excludes an additional 50bp around uncertain variants or regions. This padding eliminates many locations where Platinum Genomes only calls part of a complex or compound heterozygous variant.

In addition, we use the hap.py+vcfeval benchmarking tool (https://github.com/Illumina/hap.py) developed by the GA4GH Benchmarking Team and implemented on precisionFDA website (https://precision.fda.gov/apps/app-F187Zbj0qXjb85Yq2B6P61zb) for use in the precisionFDA “Truth Challenge”. We modified the tool used in the challenge to stratify performance by additional bed files available at https://github.com/ga4gh/benchmarking-tools/tree/master/resources/stratification-bed-files, including bed files of “easier” regions and bed files encompassing complex and compound heterozygous variants. The results of these comparisons, as well as the pipelines used to generate the calls are shared in a Note on precisionFDA (https://precision.fda.gov/notes/300-giab-example-comparisons-v3-3-2).

The callsets with lowest FP and FN rates for SNPs or indels from the precisionFDA “Truth Challenge” were compared to the v3.3.2 high-confidence calls for HG001. From each comparison result, 10 putative FPs or FNs were randomly selected for manual inspection to assess whether they were, in fact, errors in each callset.

### Integration with only Illumina and 10x Genomics WGS

To assess the impact of using fewer datasets on the resulting high-confidence vcf and bed files, we performed integration for chromosome 1 in GRCh37 using only Illumina 300x 2×150bp WGS and 10x Genomics Chromium data for HG001. We compared these calls to Platinum Genomes as we describe above for v3.3.2 calls, and we manually inspected all differences with Platinum Genomes that were not in v3.3.2.

### Differences between old and new integration methods

The new integration methods differ from the previous GIAB calls (v2.18 and v2.19) in several ways, both in the data used and the integration process and heuristics:

1. Only newer datasets were used, which were generated from the NIST RM 8398 batch of DNA (except for 10x Genomics, which used longer DNA from cells).
2. Mapping and variant calling algorithms designed specifically for each technology were used to generate sensitive variant callsets where possible: novoalign + GATK-haplotypecaller and freebayes for Illumina, the vcfBeta file from the standard Complete Genomics pipeline, tmap+TVC for Ion exome, and Lifescope+GATK-HC for SOLID. This is intended to minimize bias towards any particular bioinformatics toolchain.
3. Instead of forcing GATK to call genotypes at candidate variants in the bam files from each technology, we generate sensitive variant call sets and a bed file that describes the regions that were callable from each dataset. For Illumina GATK, we used the GATK-HC gVCF output to find regions with GQ>60. For Illumina freebayes, we used GATK callable loci to find regions with at least 20 reads with MQ >= 20 and with coverage less than 2x the median. For Complete Genomics, we used the callable regions defined by the vcfBeta file and excluded +-50bp around any no-called or half-called variant. For Ion, we intersected the exome targeted regions with the output of GATK CallableLoci for the bam file (requiring at least 20 reads with MQ >= 20). Due to the shorter reads and low coverage for SOLID, it was only used to confirm variants, so no regions were considered callable.
4. A new file with putative structural variants was used to exclude potential errors around SVs. For HG001, these were SVs derived from multiple PacBio callers (ftp://ftp-trace.ncbi.nlm.nih.gov/giab/ftp/data/NA12878/NA12878_PacBio_MtSinai/) and multiple integrated illumina callers using MetaSV (ftp://ftp-trace.ncbi.nlm.nih.gov/giab/ftp/technical/svclassify_Manuscript/Supplementary_Information/metasv_trio_validation/). These make up a significantly smaller fraction of the calls and genome (∼4.5%) than the previous bed, which was a union of all dbVar calls for HG001 (∼10%). For the AJ Trio, the union of >10 submitted SV callsets from Illumina, PacBio, BioNano, and Complete Genomics from all members of the trio combined was used to exclude potential SV regions. For the Chinese Trio, only Complete Genomics and GATK-HC and freebayes calls >49bp and surrounding regions were excluded due to the lack of available SV callsets for this genome at this time, which may result in a greater error rate in this genome. The SV bed files for each genome are under the supplementaryFiles directory.
5. To eliminate some errors from v2.18, homopolymers >10bp in length, including those interrupted by one nucleotide different from the homopolymer, are excluded from all input callsets except PCR-free GATK HaplotypeCaller callsets. For PCR-free GATK HaplotypeCaller callsets, we only include sites with confident genotype calls where HaplotypeCaller has ensured sufficient reads entirely encompass the repeat.
6. A bug that caused nearby variants to be missed in v2.19 is fixed in the new calls.
7. The new vcf contains variants outside the high-confidence bed file. This enables more robust comparison of complex variants or nearby variants that are near the boundary of the bed file. It also allows the user to evaluate concordance outside the high-confidence regions, but these concordance metrics should be interpreted with great care.
8. We now supply global phasing information from pedigree-based calls for HG001, trio-based phasing for the AJ son, and local phasing information from GATK-HC for the other genomes.
9. We use phased reads to make variant calls from 10x Genomics, conservatively requiring at least 6 reads from both haplotypes, coverage less than 2 times the median on each haplotype, and clear support for either the reference allele or variant allele in each haplotype.

## Supplementary Information

Supplementary Table 1: Detailed manual curation results of putative FPs and FNs for GIAB v3.3.2 vs. Platinum Genomes and benchmarking of BWA-GATK vs. GIAB and PG

### Outline of v3.3.2 integration script process

1. import input files
2. Preprocess vcf and bed files (preprocess_combine_vcfs.pl)
  a. unzip vcf, keep first 10 columns, removes hom ref lines, removes chr prefix, breaks into “primitive” SNPs and indels, and sorts
  b. Change sample name in vcf (column 10 of header) to make union in correct order
  c. Remove duplicate vcf lines and remove site +- 50bp from callable bed
  d. Keep only CALLABLE lines from bed if callableloci output or all lines if only 3 columns, and remove duplicate lines bed if applicable
  e. Subtract bed files listed in last columns of callsettable from each callset if the column is a 1 for each callset
  f. Add 4^th^ column to bed with DS_ “callsetname”_callable, which is added using vcfannotate to the union vcf below
3. Create union vcf with a column for each individual vcf containing the genotype call and genotype quality from the vcf (or .’s if the vcf doesn’t contain a position)
4. Annotate union vcf with each of the callable bed files in the “callable” INFO field
5. Run Integration script with “callable” annotations to get training calls for one class filtering (VcfClassifyUsingFilters_v3.3.pl)
  a. Find sites where at least 2 technologies have the same variant and genotype call, no callsets have a discordant call, and at least one callset is inside the callable bed file
6. Intersect calls found in 2 technologies with calls from each callset to give training variants to one class filter
7. Run one class filter on each callset (RunOneClassFilter.pl)
  a. One class filtering (VcfOneClassFiltering_v3.3.pl)
    i. Build an array containing the values from each annotation for 40000 random rows from the training vcf, separately for SNPs and non-SNPs
    ii. Step 2: For each annotation, sort the values from smallest to largest and find the appropriate cut-off(s)
      1. Find cut-off values for this annotation (Exclude extreme values less than or greater than 5%/(number of annotations)/(number of tails excluded))
    iii. finally, create a bed file using the vcf records that fall outside the cut-offs from Step 2 or are already filtered or GQ<20
      1. add 50bp padding to either side of filtered vcf rows
8. Annotate union vcf with each of the filtering bed files in the “oneclassfilt” INFO field
9. Run final pass of integration to find high-confidence variants using callable bed and filtering bed for arbitration (VcfClassifyUsingFilters_v3.3.pl)
  a. first, check if all genotypes that are callable agree
  b. Next, also check filtering information (To be counted, the call must satisfy: is this callset’s GT the same as the first callable callset’s GT, is this one callable, is it not filtered, is GQ>=20, and (is it not (filtered in another callset for this dataset and homozygous reference)))
    i. If all genotypes that were callable agreed, then make sure at least one is not filtered to be high confidence
    ii. If all genotypes that were callable did not agree, then see if filtering information helps to arbitrate and determine which callset to trust
    iii. Otherwise, all callsets were not callable or had low GQ, so get the first genotype of a callset that is variant
  c. Sum DP, GQ, and AD from all datasets supporting the output genotype call
  d. Filter sites incorrectly calling hets when another callset has a homozygous variant call with GQ>70 (fixes occasional erroneous calls from CG and freebayes)
  e. Filter sites that have allele balance sum < 0.2 or > 0.8
  f. Filter calls from CG if no other callsets have a variant at the position
10. Create high-confidence bed file
  a. Find regions that are callable in at least 1 callset
  b. Exclude low confidence variant positions with 50bp padding
  c. Compare high confidence variants to themselves to exclude any variants that are overlapping or represented incorrectly
  d. Exclude reference N positions
  e. For GRCh38, remove modeled centromere and heterochromatin regions
  f. For HG001/NA12878, remove ∼1Mbp somatic deletion on chr22 identified in ^6^

### Detailed manual curation results

#### Illumina Platinum Genomes

To understand the reasons for the 108 differences remaining between PG and v3.3.2 in HG001, we manually curated reads from 300x PCR-free Illumina, haplotype-separated 10x Genomics, moleculo assembled long reads, and raw and error-corrected PacBio, as well as evaluating the evidence in the Complete Genomics, SOLiD, and Ion vcf files. We found that these sites fell in several categories, and each site is assigned one of these categories and explained in detail in Supplementary File 1. (a) 23 of the PG-only calls are partial or incorrect variant calls that are at the same position or near 23 accurate v3.3.2-only calls. These are generally tandem repeats where PG calls only one half of a compound heterozygous variant (e.g., Fig. 2) or PG calls the incorrect length of an insertion or deletion. (b) 18 additional PG-only sites appear likely to be false positives in PG, mostly due to either mapping errors within a locally duplicated sequence or a systematic sequencing error that inherits properly due to its association with a nearby true variant. (c) 14 of the NIST-only sites appear very likely to be true but are missing from PG, presumably because all of the input callsets to PG missed them; 9 of these were in homopolymers or tandem repeats. (d) 3 of the NIST-only sites appear likely to be cell-line somatic de novo mutations, since they were in multiple callsets at abnormally low allele fraction, usually around 30%. These would generally not be expected to inherit properly, as discussed in the PG paper.^6^ We assign the “alleleimbalance” filter to heterozygous variants with <20% or >80% allele fraction to remove most of the low fraction variants from our high-confidence regions, but somatic mutations with moderate allele fraction remain. (e) 5 of the PG-only sites appear to be FNs in v3.3.2 due to missing variants in the CG vcf, and 1 of the v3.3.2-only variants appears to be a FP in v3.3.2 for unclear reasons. (f) 2 of the PG-only variants and 6 of the v3.3.2-only variants were unclear either due to the possible presence of a structural variant or a long repeat, and often variants were at low fractions, so these regions should probably be excluded from the high-confidence bed file. (g) Finally, 13 of the PG-only variants fell in a likely ∼1 Mbp somatic deletion in HG001 described in Fig. 3c in the PG paper.^6^ Because variant calls in this region are likely to be unreliable and batches of cells may vary in the fraction of cells with the deletion, we have excluded 22:22300000-23300000 from the final GRCh37 HG001 v3.3.2 high-confidence bed file and the corresponding chr22:21945628-22957826 from the final GRCh38 HG001 v3.3.2 high-confidence bed file. This removed most of the questionable calls in v3.3.2.

### Phasing information is provided but should be interpreted with care

The primary goal of our current integration methods is to provide high-confidence small variant and homozygous reference genotypes for benchmarking variant accuracy rather the phasing accuracy. However, as a preliminary resource to the community, we provide local phasing information for all v3.3.2 callsets and family-based phasing for HG001 and HG002. All of our v3.3.2 callsets use local phasing information from one GATK HaplotypeCaller high-coverage Illumina callset.

Pedigree-based phasing can provide global phasing of many paternal and maternal alleles. Platinum Genomes and Real Time Genomics have used the 17-member pedigree of HG001 to phase variants in HG001. Similarly, we have taken phasing information from a trio analysis of HG002, HG003, and HG004 to phase paternal and maternal alleles in HG002. Our v3.3.2 HG001 callset supplies phasing information in the priority (1) Platinum Genomes, (2) Real Time Genomics, and (3) GATK HaplotypeCaller. The Phase Set field (PS) specifies PATMAT for alleles phased by the pedigree methods in the order paternal|maternal, or takes the PID from GATK HaplotypeCaller. For HG001 GRCh37, 99.0% of high-confidence calls are phased by the Platinum Genome or RTG pedigree analyses, and 99.5% are phased by the pedigree, GATKHC, or are homozygous variant. For HG002 GRCh37, 87.0% of high-confidence calls are phased by the trio, and 89.5% of calls are phased by the trio, GATKHC, or are homozygous variant. Calls with phasing from the pedigree also have the additional level of evidence that they were inherited as expected in the pedigree. Further work is needed to evaluate the accuracy of phasing methods and integrate results from different phasing methods, including from linked reads and long reads. Therefore, if one uses these phasing results to benchmark phasing, we recommend doing so cautiously.

### GRCh38 calls are less comprehensive than GRCh37

We formed high-confidence calls for all genomes with respect to GRCh38 in a similar way to GRCh37, except that Complete Genomics, Ion exome, and SOLiD data were not mapped directly to GRCh38. Instead, vcf and bed files for these 3 datasets were converted to GRCh38 in a conservative way using GenomeWarp. All Illumina and 10x datasets were mapped directly to GRCh38. Across all genomes, total numbers of high-confidence SNPs were 1% to 8% lower than in GRCh37, and total numbers of high-confidence indels were 0% to 3% higher than in GRCh37 (detailed HG001 statistics in Table 1). High-confidence regions cover 82.0 to 85.4% of non-N bases in GRCh38 vs 87.4 to 90.8% of non-N bases in GRCh37.

To assess the accuracy of our v3.3.2 GRCh38 high-confidence calls, we compared them to the Platinum Genomes GRCh38 calls for HG001. There were 180 calls in Platinum Genomes and not 3.3.2, and 94 calls in 3.3.2 and not in PG, which is 2 to 3 times higher than for GRCh37. We manually inspected 10 random calls only in v3.3.2 and 10 only in PG to understand the source of these differences. We found that 9 of the 10 calls only in v3.3.2 were clearly supported by the data (most were missed by PG or mis-called by PG). 1 of the 10 was a complex, compound heterozygous deletion that was mis-called both in v3.3.2 and in PG. Of the 10 PG-only calls we manually inspected, 5 were inaccurate variant calls in PG near a true variant that was correct in v3.3.2, 2 were true variants missed by v3.3.2 for unclear reasons, and 3 were in difficult regions where the true genotype was unclear. 2 of these unclear variants were in 2 clusters of several PG-only SNPs on chr17 and chr19, which were regions fixed in GRCh38 and only in fix patches in GRCh37. Future work will be needed to understand accuracy of calls in these challenging regions.

### Integrating only Illumina and 10x Genomics increases high-confidence indels and errors in high-confidence regions

Some of the whole genome sequencing technologies we integrated are no longer available or in common use. Two broadly available whole genome sequencing technologies are Illumina paired end sequencing and 10x Genomics. Although 10x uses Illumina chemistries for sequencing, our analysis methods are substantially different for 10x since we use their phased haplotype-separated reads. We formed high-confidence calls for chromosome 1 by integrating only 300x Illumina WGS and 10x Genomics, using the same process and parameters for our GRCh37 HG001 v3.3.2 high-confidence calls.

Some differences exist between the Illumina-10x integrated calls and the v3.3.2 calls. The Illumina-10x and v3.3.2 high-confidence bed files are almost the same size (<0.1% different). The Illumina-10x high-confidence calls contained 1.1% fewer SNPs and 12.6% more indels than v3.3.2 in their high-confidence regions. The increase in number of indels may result from fewer sites with conflicting evidence, where, for example, Illumina has a confident indel call but another dataset has a confident homozygous reference call.

To evaluate the accuracy of the Illumina-10x integrated calls, we compared the results to Platinum Genomes, similar to how we compared v3.3.2 to Platinum Genomes above. The total number of variants matching PG were similar when comparing Illumina-10x or v3.3.2 to PG, but Illumina-10x had 1.3% fewer SNPs and 6.5% more indels matching PG. When comparing Illumina-10x integrated calls to PG, there were 10 extra PG variants and 9 extra integrated variants in the bed files (vs. 4 extra PG variants and 5 extra integrated variants when comparing v3.3.2 to PG). Since there were about 2 times more differences with PG in the Illumina-10x calls in chr1, we manually inspected all of these differences, and we found that most of the differences not in v3.3.2 were errors in the Illumina-10x integrated calls, mostly around complex variants and one large FP deletion (see Supplementary Table x for details). After manual inspection, v3.3.2 differences with PG contained only 1 erroneous extra variant and 1 unclear missing variant in chr1. After manual inspection, Illumina-10x differences with PG contained 2 partial calls of complex variants in Illumina-10x (resulting in 2 extra variants and 4 missing variants), 1 missed complex variant in Illumina-10x (resulting in 2 missing variants), 1 erroneous large deletion in Illumina-10x, and 1 unclear missing variant.

Therefore, with our current integration methods, including only Illumina and 10x whole genome data appears to result in a larger number of high-confidence indel calls but a roughly 5x increased error rate compared to integrating 4 different whole genome and 1 exome sequencing technologies. Improvements in the integration methods and calling methods may help correct many of the errors produced when using only Illumina and 10x, and incorporating long reads would also likely help.

### Performance metrics from comparison to different GIAB genomes are similar but not identical

To assess differences between genomes, we have also compared results from a single mapping and variant calling method on downsampled 50x Illumina PCR-free WGS to our high-confidence calls for all five genomes. For this analysis, we used the Sentieon haplotyper pipeline available on PrecisionFDA because the pipeline was made publicly runnable as an applet during the recent PrecisionFDA Challenge, and it is designed to mimic the widely used bwa mem and GATK HaplotypeCaller methods. 50x 150bp PCR-free Illumina data was used for all genomes except HG005, since only 250bp PCR-free Illumina data (also approximately 50x) was available for HG005.

Supplementary Table 2 shows the performance metrics obtained when comparing to each genome, stratifying SNPs vs indels and all high-confidence regions vs. all high-confidence regions minus all difficult region bed files developed by the GA4GH Benchmarking Team (including homopolymers, tandem repeats, and difficult to map regions). Indel recall and precision are similar across the 5 genomes, ranging from 99.02% to 99.26% and from 99.25% to 99.36%, respectively, in all high-confidence regions. For indels, when removing all variants in repetitive regions, the FP rate was similar for all genomes, but the FN rate for HG005 was over 2 times higher than for any of the other genomes, for unclear reasons. Similarly, for SNPs, when removing all variants in repetitive regions, the FP rate was similar for all genomes, but the FN rate for HG005 was over 7 times higher than for any of the other genomes. Interestingly, for SNPs in all high-confidence regions, the FP rate for HG005 is ∼2 times higher but the FN rate is ∼2 times lower. Future work will be needed to determine whether the differences for HG005 is caused by the longer 250bp Illumina reads used with Sentieon, the different types of data that went into forming the benchmark calls, or the sample’s Asian ancestry. The fraction of variants not assessed because they are outside the high-confidence regions varies by 4 % or less between genomes, though HG001 has the lowest fraction not assessed in general, suggesting its characterization may be the most comprehensive. Overall, 16 to 20 % of SNPs and 45 to 49% of indels fall outside the v3.3.2 high-confidence regions, which highlights the need for more comprehensive high-confidence calls and regions.

**Supplementary Table 2:** Performance metrics from comparing a single mapping and variant calling method to our v3.3.2 GRCh37 high-confidence calls for each genome. Statistics are separated for SNPs and indels and for all high-confidence regions and high-confidence regions minus “all difficult regions” (homopolymers, tandem repeats, and difficult to map regions). 50x 150bp PCR-free Illumina data was used for all genomes except HG005, since only 250bp PCR-free Illumina data was available for HG005.

| Genome | Type | Subset | 100% -<br>recall | 100% -<br>precision | Recall | Precision | Fraction of<br>calls outside<br>high-conf bed |
| --- | --- | --- | --- | --- | --- | --- | --- |
| HG001 | SNP | all | 0.0277 | 0.1274 | 0.9997 | 0.9987 | 0.1653 |
| HG002 | SNP | all | 0.0664 | 0.1342 | 0.9993 | 0.9987 | 0.1910 |
| HG003 | SNP | all | 0.0625 | 0.1489 | 0.9994 | 0.9985 | 0.1967 |
| HG004 | SNP | all | 0.0633 | 0.1592 | 0.9994 | 0.9984 | 0.1975 |
| HG005 | SNP | all | 0.1175 | 0.0870 | 0.9988 | 0.9991 | 0.1834 |
| HG001 | SNP | notinalldifficultregions | 0.0096 | 0.0783 | 0.9999 | 0.9992 | 0.0491 |
| HG002 | SNP | notinalldifficultregions | 0.0102 | 0.0576 | 0.9999 | 0.9994 | 0.0864 |
| HG003 | SNP | notinalldifficultregions | 0.0128 | 0.0819 | 0.9999 | 0.9992 | 0.0864 |
| HG004 | SNP | notinalldifficultregions | 0.0102 | 0.0860 | 0.9999 | 0.9991 | 0.0854 |
| HG005 | SNP | notinalldifficultregions | 0.0931 | 0.0541 | 0.9991 | 0.9995 | 0.0664 |
| HG001 | INDEL | all | 0.8354 | 0.7458 | 0.9916 | 0.9925 | 0.4485 |
| HG002 | INDEL | all | 0.8271 | 0.7016 | 0.9917 | 0.9930 | 0.4547 |
| HG003 | INDEL | all | 0.7546 | 0.6523 | 0.9925 | 0.9935 | 0.4632 |
| HG004 | INDEL | all | 0.7345 | 0.6390 | 0.9927 | 0.9936 | 0.4592 |
| HG005 | INDEL | all | 0.9840 | 0.7418 | 0.9902 | 0.9926 | 0.4850 |
| HG001 | INDEL | notinalldifficultregions | 0.0551 | 0.1475 | 0.9994 | 0.9985 | 0.1927 |
| HG002 | INDEL | notinalldifficultregions | 0.0497 | 0.0893 | 0.9995 | 0.9991 | 0.2208 |
| HG003 | INDEL | notinalldifficultregions | 0.0508 | 0.1627 | 0.9995 | 0.9984 | 0.2229 |
| HG004 | INDEL | notinalldifficultregions | 0.0496 | 0.1307 | 0.9995 | 0.9987 | 0.2190 |
| HG005 | INDEL | notinalldifficultregions | 0.1182 | 0.1535 | 0.9988 | 0.9985 | 0.2049 |

### Performance metrics can vary significantly when comparing to different benchmark callsets

Errors and partial calls in the benchmark callset can cause precision and recall to appear higher or lower than they should, and previous work has proposed methods to account for these errors in the uncertainty of precision and recall values.^20^ It is generally difficult to estimate the error rates of benchmark callsets, because they are very low and truth is not known. We approximate error rates above through comparison to an orthogonal benchmark in Platinum Genomes along with manual inspection of the differences and trio mendelian analysis.

Simplistic comparison methods that do not account for different representations of complex variants can cause methods that represent variants similarly to the benchmark to appear better than methods that represent variants in a different way.^21,22^ The Global Alliance for Genomics and Health Benchmarking Team has developed sophisticated methods that account for these differences as long as the complex variant is fully called correctly in both the benchmark and test callsets.^8^ Differences in representation can make up a significant fraction of putative FPs and FNs, so it is important to use these tools.

Another consideration is that benchmark callsets and regions tend to be biased towards easier variants and easier genome contexts. This bias generally causes sensitivity and precision estimates to appear better than they would if the benchmark included all variants in the genome.

A striking example of the importance of understanding limitations of benchmark callsets is in regions of the genome that are homologous to the often used hs37d5 decoy sequence for GRCh37 from the 1000 Genomes Project.^23^ The decoy was created to capture reads that are from contigs that are not in the GRCh37 reference assembly, which otherwise can cause FPs. Our current high-confidence regions exclude regions homologous to the decoy except when 10x data has normal coverage of both haplotypes in these regions and each haplotype has near 100% of the reads supporting either reference or alternate. One possible disadvantage of using the decoy is that it may cause FNs if the decoy sequence is not a homologous duplication in the genome but is actually an alternate haplotype in the region homologous to it. Since 10x includes the decoy when mapping reads, if the decoy is an alternate haplotype then 10x should only have reads from one haplotype in the region homologous to the decoy, and it would not meet our criteria for high-confidence. Therefore, we include 10x calls in our high-confidence regions only if there is normal coverage on both haplotypes, and we exclude all other callsets, which do not have haplotype information.

We compared recall, precision, and fraction not assessed for the decoy-associated regions when comparing bwa-GATK calls without the decoy (bwa-GATK-nodecoy) to our v3.3.2, v2.18, and PG high-confidence calls. The precision for bwa-GATK-nodecoy is 67% vs. v3.3.2, whereas it is 91% vs. v2.18 and 93% vs. PG. The much higher precision vs v2.18 or PG is a result of many of the decoy-associated variants being excluded from the high-confidence regions of v2.18 and PG. For v2.18, these variants are excluded due conflicts between datasets using the decoy and those that do not, and for PG these variants are excluded because they do not inherit properly. Assessing these variants is important because they make up 7123 of the 16452 FPs (43%) vs. v3.3.2 for this callset, even though variants in decoy-associated regions only make up 0.5% of TPs.

The sensitivity of bwa-GATK-nodecoy and bwa-GATK-decoy in decoy-associated regions is >99% vs v2.18 and v3.3.2, but when compared to PG the sensitivity of bwa-GATK-nodecoy is 93% and bwa-GATK-decoy is 92%. In this way, PG appears to be better at discriminating FNs. When manually examining these, they generally are in regions with a high density of variants and often with low mappability, which makes them difficult to call accurately, at times even when they inherit as expected.

Interestingly, 91%, 92%, and 78% of variants from bwa-GATK-nodecoy in the decoy-associated regions are outside the high-confidence regions of v2.18, v3.3.2, and PG, respectively. Therefore, PG is assessing the most variants, and it appears to find the most FNs, but v3.3.2 finds the highest proportion of FPs despite assessing the lowest number of variants. This strength of v3.3.2 likely results from including more regions prone to FPs when not using the decoy.

We also manually inspected a random subset of putative FP deletions >15bp from bwa-GATK (including the decoy when mapping). When compared to PG, the precision for these was 75%, whereas when compared to v3.3.2 the precision was 97%. When inspecting 10 random FPs compared to v3.3.2, we found 3 were likely errors in bwa-GATK not near a true variant, 4 were unclear, and 3 were true deletions, all >50bp. These deletions >50bp are not excluded by the v3.3.2 bed because only 50bp on either side of the start of an uncertain deletion are excluded from the bed, which will be fixed in future versions. When inspecting 10 random FPs compared to PG, we found 1 was a likely error in bwa-GATK not near a true variant, 1 was unclear though the true variant is not in PG, and 8 were accurate deletions that were only partially excluded by the PG bed. In this case, neither benchmark is perfect, but because many of the putative FPs vs PG are not actually errors, the precision of 75% is likely a significant underestimate of the true precision of bwa-GATK inside the PG bed file. Note, however, that the true precision of bwa-GATK is unknown outside the v3.3.2 and PG bed files.

Similarly, we manually inspected a random subset of putative FP indels that were in or near compound heterozygous variants from bwa-GATK. We used compound heterozygous indel variants from the RTG-phased pedigree calls^7^ to minimize bias towards v3.3.2 or PG. When compared to PG, the precision for these was 93%, whereas when compared to v3.3.2 the precision was 95%. When inspecting 10 random GATK FPs compared to v3.3.2, we found 9 were likely errors in bwa-GATK mis-genotyping the true compound heterozygous event, and 1 was a likely mis-genotype in v3.3.2. When inspecting 10 random GATK FPs compared to PG, we found 4 were likely errors in bwa-GATK mis-genotyping the true compound heterozygous event, and 6 were accurate compound heterozygous events that were partially called or missed by PG. In this case, the v3.3.2 appears more useful than PG at finding real FPs, and because many of the putative FPs vs PG are not actually errors, the precision of 93% is likely an underestimate of the true precision of bwa-GATK inside the PG bed file. However, again, the true precision of bwa-GATK is unknown outside the v3.3.2 and PG bed files.

A strength of pedigree-based analyses like Platinum Genomes and Real Time Genomics is their ability to assess sensitivity for some more difficult variants. For example, when bwa-GATK is compared to PG the sensitivity to SNPs in RefSeq coding regions is 97.98%, whereas when bwa-GATK is compared to v3.3.2 the sensitivity to SNPs in RefSeq coding regions is 99.96% (a FN rate estimate 62 times lower). To determine the accuracy of the extra variants in PG, we manually inspected 10 random FNs in bwa-GATK relative to PG. We found that all of these were in difficult-to-map regions. 8 of the 10 had evidence in most of the long or linked read technologies (PacBio, 10x, and Moleculo), though some had low mapping quality or high coverage even in long and linked reads, potentially related to duplications and mapping errors. 2 of the 10 did not have evidence in long or linked read technologies and were in low mapping quality and/or discordantly mapped reads in short read technologies, so may be errors related to mis-mapping or local duplications. While some putative FNs may be inaccurately called variants in PG, PG does help identify more questionable variants in difficult-to-map regions than v3.3.2, since v3.3.2 excludes most of these variants from its high-confidence regions.

### Limitations of v3.3.2 Benchmark Calls

While our newest version of high-confidence calls is more comprehensive and accurate than previous versions, it is essential to understand its limitations as well. Benchmarking statistics obtained upon comparison to our calls likely will not accurately reflect performance metrics for all variants in the whole genome, because they are not a representative sample of all variants.

Importantly, the most difficult regions of the genome and types of variants are underrepresented in our high-confidence vcf and bed files. Regions difficult to map with short reads, as well as long homopolymers and tandem repeats, are generally excluded from our high-confidence regions. In addition, larger indels and structural variants are generally excluded, and some complex variants that could not be normalized are excluded. Also, some true deletions >50bp in length are not completely excluded by our bed file and may appear incorrectly to be false positives.

Currently, only the autosomes (chromosomes 1 to 22) are included in our high-confidence regions for all genomes; HG001 is the only genome for which chromosome X is characterized. We do not currently have calls for Y, MT, or any of the unlocalized or unplaced contigs or ALT loci for GRCh37 or GRCh38. Performance in these regions is likely to differ, partly because they are not necessarily diploid (e.g., X and Y in males and MT in everyone) or because individuals are often highly divergent from the reference (e.g., in the HLA genes).

Many de novo mutations are likely to be excluded from our high-confidence regions, particularly if they are in only a fraction of the cells (e.g., somatic mutations arising in the cell line)

However, we have found that some potential de novo mutations are included in our high-confidence regions; higher fraction mutations may be called heterozygous and low fraction mutations may be called homozygous reference, which can cause some putative FPs and FNs.

Regions with clusters of variants on one allele sometimes are mistakenly called high-confidence homozygous reference. These variants are sometimes missed if reads containing variants do not map to the region. For future versions, we are exploring using graph-based alignment methods and better local and global assembly methods to call these regions.

For all of these reasons, benchmarking statistics should always be examined critically. If one technology or variant caller has better overall performance metrics than another method, it does not always mean that the first method is the best methods for all types of variants, or that it better detects the variants in which one is interested (e.g., indels are often overrepresented in clinically significant variants). It is important both to understand the variant types and regions being assessed and those not being assessed by the current high-confidence set.

### Benchmarking Best Practices

To compare to GIAB high-confidence calls and obtain robust, standardized performance metrics, we strongly recommend following the best practices established by the Global Alliance for Genomics and Health Benchmarking Team for germline small variant calls.^8^ The tools developed by this team enable sophisticated comparisons of differing representations of complex variants, output of standardized performance metrics like true positive, false positive, false negative, and genotype error, comparison at different stringencies of matching (e.g., requiring genotype match, allele match, or local match), and stratification of performance metrics by variant type and genome context. These best practices can currently be followed most easily by using the hap.py toolkit on GitHub or its implementation on precisionFDA, with up-to-date guidelines available at https://github.com/ga4gh/benchmarking-tools.

